# Nonlinear transfer and temporal gain control in ON bipolar cells

**DOI:** 10.1101/514364

**Authors:** Nikhil R. Deshmukh, Michael J. Berry

**Affiliations:** Princeton Neuroscience Institute, Princeton University, Princeton, New Jersey 08544

## Abstract

The separation of visual input into discrete channels begins at the photoreceptor to bipolar cell synapse. Current models of the ON pathway describe the time-varying membrane voltage of ON bipolar cells as a linear function of light fluctuations. While this linearity holds under some visual conditions, stimulating the retina with full-field, high contrast flashes reveals a number of nonlinearities already present in the input current of ON bipolar cells. First, we show that the synaptic input to ON bipolar cells is asymmetric in response to equal flashes of opposite polarity. Next, we show that this asymmetry emerges because the responses to dark flashes increase linearly with contrast, whereas responses to bright flashes are highly rectified. We also describe how the outward current saturates in response to dark flashes of increasing duration. Furthermore, varying the inter-flash interval between a pair of high contrast flashes reveals a rapid, transient form of gain control that modulates both the amplitude and time course of the flash response. We develop a phenomenological model that captures the primary features of the ON bipolar cell response at high contrast. Finally, we discuss the implications of these nonlinearities in our understanding of how retinal circuitry shapes the visual signal.

## Introduction

Bipolar cells, the second-order neurons of the retina, are increasingly understood to be the site of important and powerful visual computations [1]. Their axon terminals can convert membrane voltage into a graded release of glutamate in a manner embodying rectification of the visual signal. As a result of this rectification, bipolar cells act as nonlinear subunits within a ganglion cell’s receptive field [2–5], thereby giving rise to forms of translation invariance in retinal motion processing [6–8]. Furthermore, synaptic depression within the axon terminal can dramatically reduce glutamate release [9–11], giving rise to a form of gain control that emphasizes motion discontinuities [8, 12].

In addition to these image transformations at its output, bipolar cells can also carry out powerful computations in their dendrites through the action of their glutamate receptors. In order to more efficiently encode increments and decrements of light intensity, the retina splits the visual signal into parallel ON and OFF pathways [13]. OFF bipolar cells have ionotropic receptors that preserve the (inverted) sign of visual signals, and both kainate- and AMPA-type receptors exhibit a strong form of desensitization with different kinetics [14, 15]. ON bipolars have a metabotropic receptor, mGluR6, that inverts the sign of their glutamate signals to produce depolarizing responses to increases in light intensity [16]. Activation of mGluR6 by binding of glutamate triggers a second messenger cascade that ultimately closes the cation channel TRPM1 [17–20], although the full biochemical pathway has not been delineated. Because ON bipolars use this signaling cascade, they have the potential to carry out even more sophisticated computations within their dendrites than OFF bipolars.

Indeed, at low light level rod bipolar cells have a threshold nonlinearity that helps improve signal-to-noise ratio for the integration of single-photon events generated by rod photoreceptors [21]. This nonlinearity results from saturation in the mGluR6 cascade [22]. At higher light levels, ON bipolar cells exhibit saturation for both positive and negative contrasts [23, 24]. The mGluR6 cascade exhibits desensitization via feedback from intracellular calcium on a timescale of ~0.8 sec [25] as well as potentiation via cGMP [26]. At shorter timescales, however, the dynamics of the cascade are not as well characterized. Another approach to studying the transformation of light to ON bipolar cell activity has ignored this biophysical complexity. Instead, the response characteristics were elucidated using reverse correlation to spatiotemporal white noise stimulation. Under these conditions, most of the fluctuations in membrane voltage were captured by a linear model [27–30] with bandpass temporal kernels [28, 31]. This kernel can adapt as a function of the mean membrane voltage due to the action of voltage-gated K+ channels [32].

We seek to reconcile these different views and produce a unified model of ON bipolar cell function at the level of its synaptic input current. Our approach has been to study the synaptic currents into ON bipolar cells using voltage clamp measurements in the retinal slice preparation. We varied the contrast and duration of bright and dark flashes of light to probe both the linear and saturating regimes. We find that the synaptic input to ON bipolar cells already contains multiple sources of significant nonlinearity that are present for moderate contrast and short duration flashes of light. There are two sources of saturation giving rise to responses that are asymmetric for bright versus dark flashes. Input currents have a biphasic temporal kernel in which the overshoot increases strongly as a function of contrast. In addition, we found a gain control mechanism acting on the same time scale as the immediate light response. We describe a unified computational model that captures these nonlinearities in the ON bipolar cell light response. This model includes saturation at the photoreceptor-to-bipolar cell synapse, temporal gain control within the mGluR6 cascade, and saturation in the opening of the TRPM1 channel.

## Materials and Methods

### Preparation

Experiments were performed on larval tiger salamanders (Charles Sullivan, Nashville, TN) kept at 16°C on a 12-hr light–dark cycle. Care and euthanasia of the animals were carried out in accordance with procedures approved by Princeton University Animal Care and Use Committee. After being dark-adapted for 2 hours, salamanders were rapidly decapitated and the head and spinal cord pithed under dim illumination; all subsequent steps were performed under infrared illumination. The eyes were removed and hemisected, and the cornea, iris, lens, and vitreous humor were removed. The retina was embedded in low melting temperature agarose (Sigma) made with HEPES-buffered AMES medium and sliced into 250 um thick slices on a vibrating microtome (Leica) at 4°C.

### Electrophysiology

All recordings were made at room temperature in bicarbonate-buffered Ringers solution containing (in mM): 110 NaCl, 22 NaHCO_3_, 2.5 KCl, 1.6 MgCl_2_, 1 CaCl_2_, and 10 glucose, equilibrated with 95% O_2_, 5% CO_2_, and adjusted to pH 7.35 with NaHCO_3_. Retinae were viewed under infrared illumination, and individual bipolar cells were patched using standard whole-cell patch techniques. Patch electrodes were pulled from 1.2 mm OD borosilicate glass (FHC) on a Sutter P-2000 micropipette puller. Electrode diameters were approximately 1.0 μm at the tip, and had resistances of 8-12 MΩ. All whole-cell patch recordings were performed with a Multiclamp 700B patch-clamp amplifier (Molecular Devices). Bipolar cell intracellular solution contained (in mM): 125 K-aspartate, 10 KCl, 10 HEPES, 5 EDTA, 1 CaCl_2_, 1 ATP, and 0.5 GTP adjusted to pH 7.3 with NMG-OH. Bipolar cell types were confirmed by measuring light responses, and cells with a series resistance >50 MΩ were discarded. The liquid junction potential (−9mV) was uncorrected.

### Visual stimulation

Full field flash stimuli were delivered by a 670 nm LED (SuperBrightLEDS) behind a holographic diffuser (Thorlabs) focused onto the retina from the bottom of the recording chamber. Bright and dark flashes of high contrast (±1.0 Weber contrast) and varying duration were interleaved and presented on background illumination to define the adaptational state of the retina in the photopic regime. We presented a set of 50 random binary white noise sequences, each having a duration of 5 seconds, a bandwidth of 30 Hz, and a temporal contrast of 33% (standard deviation across time divided by the mean). Of these sequences, 15 were identical and served as the test sequence for the linear model, and the remaining 35 were unique and were used to calculate the parameters of a linear-nonlinear model. The mean background light intensity of all stimuli measured at the plane of the retina was 1650 photons/L-cone/sec. Photon flux was calculated using salamander photoreceptor spectra values reported in the literature [33, 34]. The retina was continuously illuminated by a rod-suppressing 540 nm green background to isolate cone circuitry.

### Analysis

Data were acquired at a sampling rate of 20 kHz and low-passed filtered at 1 kHz using pClamp software (Axon Instruments, Foster City, CA). Subsequent analysis was performed using SciPy, an open-source scientific computing environment for the Python programming language. The linear filter in Figure 3A was calculated as in [35]. Following [36], the nonlinearity (Figure 3B and 4A) was determined by binning the predicted response into 0.5 mV bins and calculating the average measured current response for each bin.

### Computational Model

We developed a Linear-Nonlinear-Kinetic-Nonlinear (LNKN) model to predict the synaptic input current to ON bipolar cells over the entire range of visual stimuli that we tested experimentally. The stimulu*s s(t)* is the Weber contrast of the full-field luminance *Y* as a function of time:

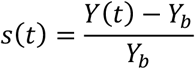

where*Y*_b_ is the background luminance.

We modeled the photoreceptor voltage *L(t)* as the convolution of the stimulus *s(t)* with a linear kernel *k(t)* as follows:

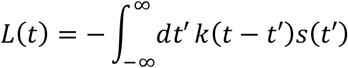

where

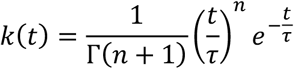

The negative sign represents the sign-inversion that produces a hyperpolarization in response to a positive contrast flash. The glutamate release at the photoreceptor *u(t)* was given by the nonlinear function, *N*_*1*_:

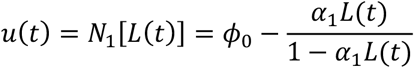

where *ϕ*_0_ represents the level of steady-state glutamate release at the background light level (*L* = 0), and *α*_1_ determines the slope of the input-output function. The units of glutamate release are truncated to the (normalized) range [0,1].

The output of the photoreceptor, *u(t),* is then passed through the kinetic block of the model, similar to [37]. This block captures the dynamics of the mGluR6 receptor to TRP channel current system by reducing all of the response elements and second messengers to three internal state variables: *R* (ready pool of signaling elements), *A* (active pool), and *X* (inactive pool). The time evolution of these internal variables is given by a Markov process:

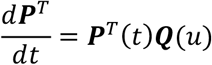

where the fractional occupancies *R*, *A*, and *I* each make up a row in the column vector **P**(*t*) and satisfy the condition *R* + *A* + *I* = 1. **Q** is a 3×3 transition matrix that determines the transitions between each of three states:

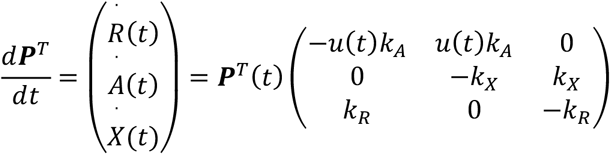

Glutamate release *u(t)* scales the rate at which a signaling element in the *R* pool transitions to the *A* pool. The solution to this system of differential equations was obtained numerically. To determine the output of the kinetic block *c(t),* we multiplied the occupancy of the active pool *A(t)* by −1 to capture the sign-inversion where the channel closes when activated and subtracted the initial value at *t* = 0, *A(0)*

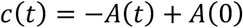

Next, to convert *c(t)* to an input current *I(t),* we applied a static nonlinearity *N*_*2*_*(t)* and scaled the response to units of pA.

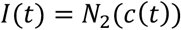

where *N* is given by:

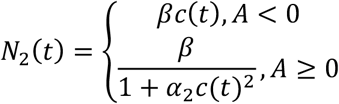

The term *β*is the scale factor from arbitrary units to pA, and *α*_2_ sets the slope of the rectification function.

The best fit parameters were determined using stochastic gradient descent, and are as follows:

| Parameters | Description | Cell 3011 |
| --- | --- | --- |
| $n$ | Exponent of linear photoreceptor kernel | 5.51 |
| $\tau$ | Time constant of linear photoreceptor kernel | 10.1 ms |
| $\phi_0$ | Constitutive glutamate release | 1.1 |
| $\alpha_1$ | Slope of photoreceptor output nonlinearity | 0.52 |
| $k_R$ | Transition rate, X→R | 0.0015/ms |
| $k_A$ | Transition rate, R→A | 0.010/ms |
| $k_X$ | Transition rate, A→X | 0.011/ms |
| $\alpha_2$ | Slope, saturating nonlinearity | 10.5 |
| $\beta$ | Scale factor, arbitrary units to pA | 31.8 pA |

## Results

Linear superposition is a powerful and important mathematical property. When it holds, the complete response of a system can be derived from well-developed mathematical methods. On the other hand, linear systems embody limited computational powers, as the input can be recomputed from the output (assuming no noise is added). Thus, a nonlinear system, while more complicated to characterize, can be said to perform more substantial processing on its inputs. To directly test the linearity of the synaptic input to ON bipolar cells, we performed voltage-clamp recordings of ON bipolar cells in a slice preparation of the tiger salamander retina in response to full-field bright and dark flashes (see Materials and Methods). ON bipolar cells were identified by their morphology and depolarizing response to bright flashes, and were clamped at their resting membrane potential as measured in current clamp (mean ± SEM = −41.2 ± 1.4 mV, n=12).

We tested whether the synaptic input to these cells was linear at high contrast by comparing the amplitudes of the peak postsynaptic current in response to 50 ms full field flashes of ±1.0 Weber contrast. To facilitate this comparison visually, the mean dark flash response (blue trace) was inverted and superimposed on the mean bright flash response (red trace), as shown in Figure 1. The response to a strong flash is biphasic, with a sharp rise to peak and a subsequent overshoot in the opposite direction. In all ON bipolar cells recorded, the amplitude of the peak outward current and the peak inward overshoot in response to a dark flash was greater than the amplitudes of the corresponding peak and overshoot in response to a bright flash.

**Figure 1.**
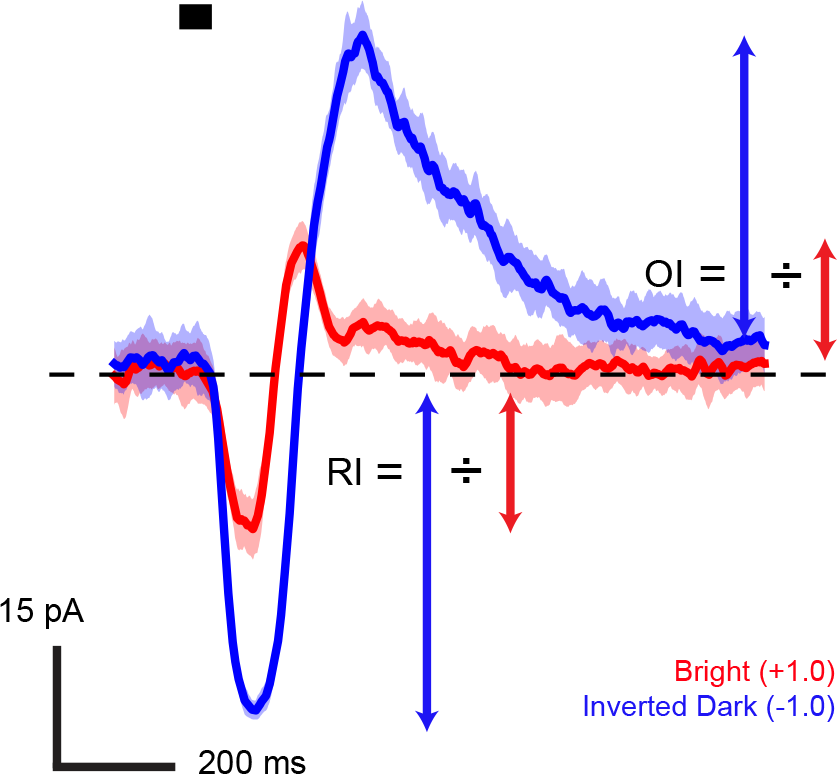
Comparison of the response to bright and dark flashes. Measured input current for an example ON bipolar current in response to 100% contrast bright (red trace) and dark flashes (blue trace). The black bar represents the duration of the 50 ms full-field stimulus. The solid traces represent the mean response and the light shaded regions around the traces show the standard deviation of the responses across 5 trials. The rectification index (RI) and the overshoot index (OI) were used to quantify response asymmetry, as illustrated.

To quantify this asymmetry between ON and OFF responses, we defined the Rectification Index (RI) of the cell as the ratio of the dark flash response amplitude to the maximal bright flash response amplitude. Analogously, we defined the Overshoot Index (OI) as the ratio of the overshoot in response to dark and bright flashes. For our population of cells, the mean RI was 2.70 ± 0.41 and the mean OI was 2.46 ± 0.31 (mean ± SEM, n=12), indicating significant rectification of light increments compared to decrements at the cone to ON bipolar synapse (RI: p=0.008; OI: p=0.015, paired t-test).

To test whether this asymmetry persists at lower contrasts, we measured the rectification of the peak and overshoot in response to flashes of ±0.25 and ±0.5 Weber contrast. The maximal amplitude of the dark flash peak response and overshoot increased linearly as a function of contrast, whereas the corresponding peak and overshoot in response to a bright flash saturated at high contrast. This led to the increase of the mean rectification index from approximately 1 (symmetric) at 25% contrast to greater than 2.5 (highly asymmetric) at 100% contrast. The emergence of this asymmetry as a function of contrast is shown for an example cell in Figure 2A. The saturation of the inward current in response to bright flashes at 50% contrast suggests that a signaling element within the ON bipolar pathway is fully activated such that a stronger stimulus cannot activate it further.

**Figure 2.**
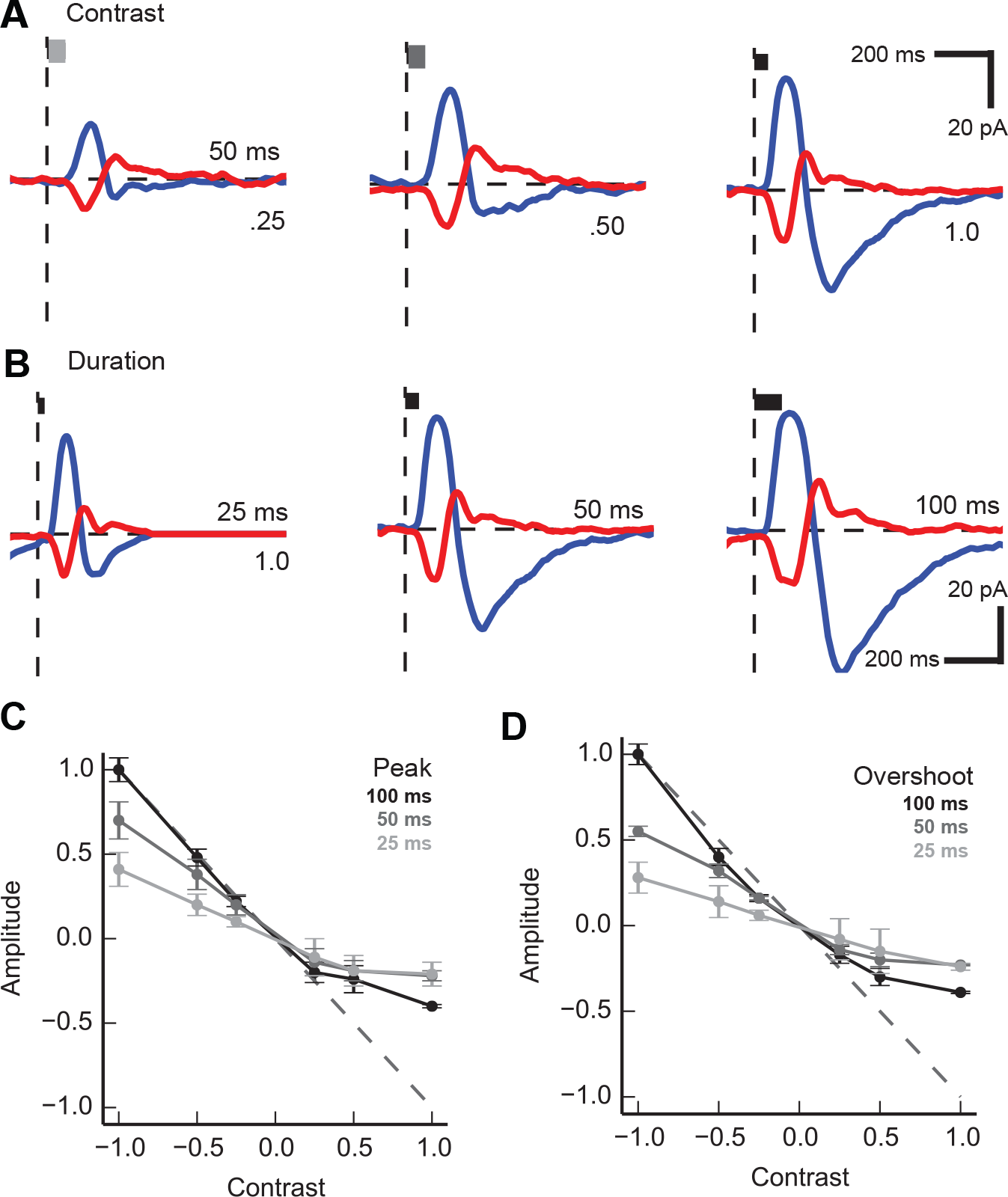
Response asymmetry between bright and dark flashes increases with contrast and flash duration. **A.** ON bipolar postsynaptic current in response to bright and dark flashes of 25%, 50% and 100% contrast. Traces represent mean response across 5 trials. **B.** ON bipolar postsynaptic current in response to flashes of 25, 50 and 100 ms duration. Traces represent mean response across 5 trials. **C.** Peak response amplitude at all contrasts and durations normalized by the peak response at 100% contrast and 100 ms duration. Dotted line indicates unity and error bars represent SEM (n=7). **D.** Peak overshoot amplitude at all contrasts and durations normalized by the peak overshoot at 100% contrast and 100 ms duration. Dotted line indicates unity and error bars represent SEM (n=7).

Next, we examined the effect of changing the flash duration on the amplitude of the peak and overshoot. This is shown for an example cell in Figure 2B. The inward current in response to a bright flash saturated as the duration of the flash increased from 25 to 100 ms. Similarly, the peak outward current in response to a dark flash also saturated over this range of flash durations. This suggests that sufficiently strong activation to produce rectification in the ON bipolar pathway can occur through temporal integration.

Figure 2C shows the contrast response curve of the peak amplitude for all ON bipolar cells recorded. The amplitude was normalized to the peak response at the highest contrast. The saturation of the ON response required flashes of sufficient contrast and duration. For 50 ms and 100 ms flashes, the slope of the response amplitude as a function of contrast was significantly shallower at positive contrasts compared to negative contrasts. At 25 ms, however, the slope was approximately constant from −1.0 to +0.5 contrast, with attenuation only at the +1.0 contrast condition.

The overshoot response normalized by the maximal overshoot (at high contrast and long duration) is shown in Figure 2D. The amplitude of the overshoot followed the same overall trend as the peak amplitude for all contrasts and durations. At low contrast and short duration, the overshoot was equally small for bright and dark flashes (compare contrasts ±1.0), consistent with a lack of rectification in this limit. However, as the flash duration was increased to 50 and 100 ms, the overshoot in response to dark flashes grew significantly larger than that for bright flashes. The similarity of the overshoot contrast response to the peak contrast response suggests that the overshoot is a consequence of the saturation of the ON bipolar pathway.

Importantly, the peak outward current in response to dark flashes shown in Figure 2B does not increase with duration of the flash. In systems theory, the principle of linear superposition states that for all linear systems, the total response produced by the sum of many stimuli presented simultaneously is equal to the sum of the individual response of the system to each stimulus alone. Testing this superposition principle provides another method to probe the linearity of bipolar cell inputs. In Figure 3, we directly compared the OFF response of a single 50 or 100 ms flash to the superposition model, in which we summed either two or four time-shifted copies of the OFF response to a 25 ms flash. For the example cell shown, the peak amplitudes of the 50 ms and 100 ms superposition model were 60 pA and 116 pA, respectively, but the measured peak outward current to either a 50 ms and 100 ms flash never exceeded 52 pA (Figure 3A). Furthermore, the superposition model does not predict the large overshoots measured in response to 50 ms and 100 ms flashes.

To examine whether the rectification of the outward current depended on contrast, we tested for linearity using the 100 ms flash superposition model at 50% and 25% contrast. For each value of the superposition model, we averaged the corresponding values of the measured response. We then normalized all the responses by the peak outward current of the measured response at 100% contrast. As shown in Figure 3B, the normalized response at 100% contrast saturates quite strongly, with the peak amplitude of the superposition model reaching more than twice the peak amplitude of the measured data. At 50% contrast, the response still saturates, but the ratio of peak amplitudes is only 1.1 for the example cell shown. At 25% contrast, the measured response equal to the superposition model, indicating linearity. Figure 3C shows the mean ratio of peak amplitudes for the 50 ms and 100 ms superposition models and measured responses for all ON bipolar cells measured. Even for the shorter 50 ms flashes, the superposition model overpredicts the actual outward current measured. This suggests that there is a limit to the maximal outward current that an ON bipolar cell can sustain, and that the cell reaches this limit in response to high contrast dark flashes. At lower contrasts, this limit is not fully reached, so the response is more linear.

**Figure 3.**
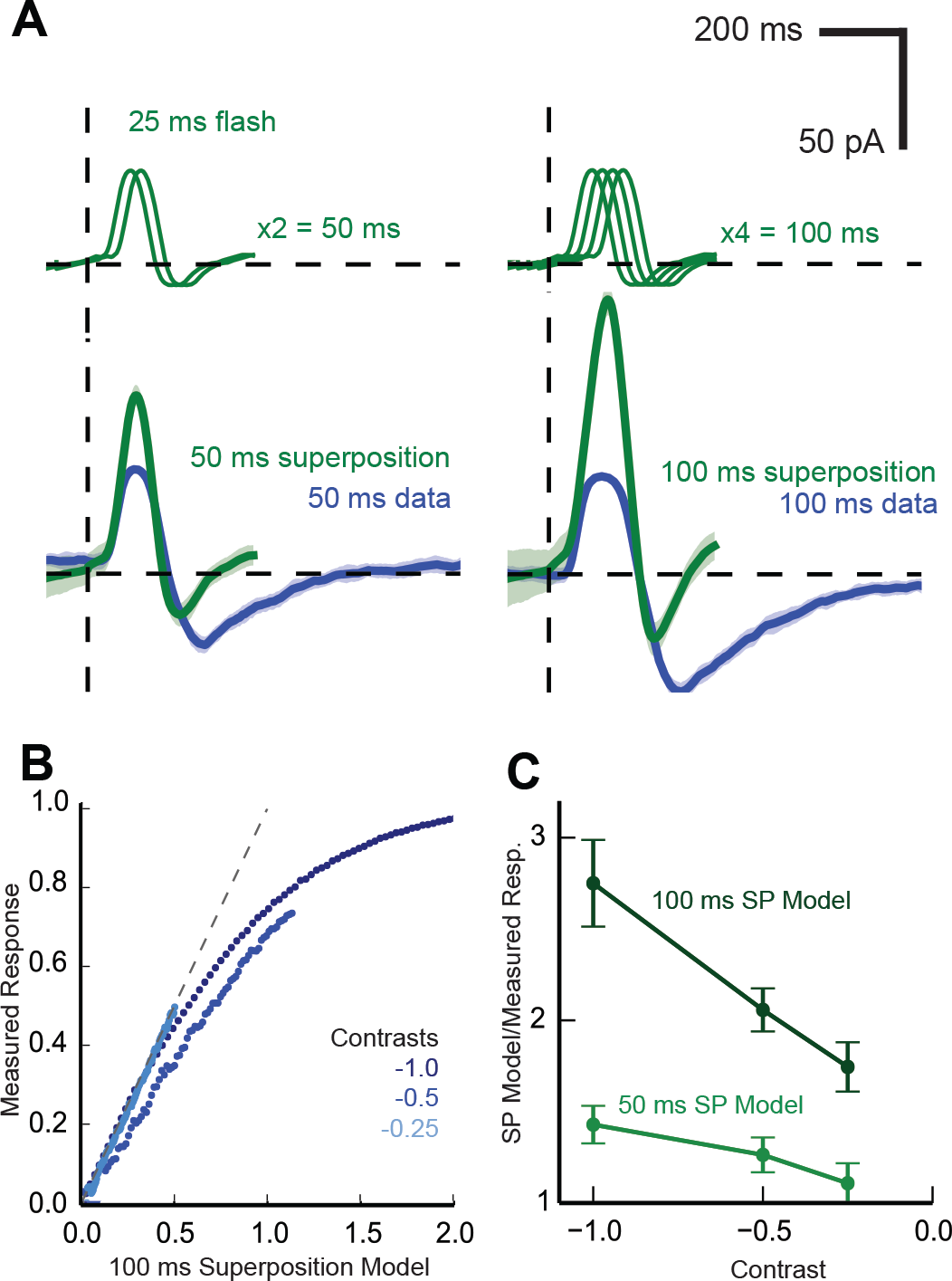
Linear superposition analysis reveals saturation of the OFF response. **A.** The sum of two or four time-shifted responses to a 25 ms flash (green) were compared with the measured response to a 50 ms and 100 ms flash, respectively (blue). Light shading represents standard deviation (5 trials). **B.** The average measured OFF response (peak outward current, normalized) to 100 ms flashes plotted against the predicted response from the superposition of 25 and 50 ms flashes for 25%, 50% and 100% contrast (shades of blue). **C.** The population average ratio of peak superposition model OFF response to measured OFF response for 50 ms and 100 ms superposition models (n=7). Error bars represent SEM.

The results from Figures 1-3 demonstrate rectification of inward and outward currents at the input to ON bipolar cells. This seems to contradict the linearity predicted by the LN models constructed in previous studies. In [28], for example, the transformation of light to salamander ON bipolar membrane voltage was characterized using low contrast (30%) white noise stimulation. The nonlinearity in this model was measured by directly comparing the predicted membrane response to measured voltage response at the ON bipolar soma. This function was found to be essentially linear. This result is consistent with our finding that there is little to no rectification at lower contrasts (Figure 2A, C). We sought to verify that the nonlinear cells we recorded were truly linear at lower contrast. We presented interleaved unique and repeated 5 sec sequences of binary white noise at 33% contrast and measured the postsynaptic current to construct an LN model. The linear filter (Figure 4A) and nonlinear function (Figure 4B) were calculated from the responses to the unique sequences as described previously in [28, 35]. The linear filter was biphasic, with a transient inward component around 100 ms, and a broader outward component between 150 and 300 ms. The measured nonlinearity approximated a straight line and did not saturate, demonstrating a lack of rectification at 33% contrast. This LN model was then applied to the repeated stimulus sequence and compared to the average measured response (Figure 4C). While the model does not exactly match the measured response, the normalized root mean squared error between model and measured current was 18.1 ± 2.0% (±SEM, n=7), indicating that the model accounted for more than 80% of the structure of the light response.

**Figure 4.**
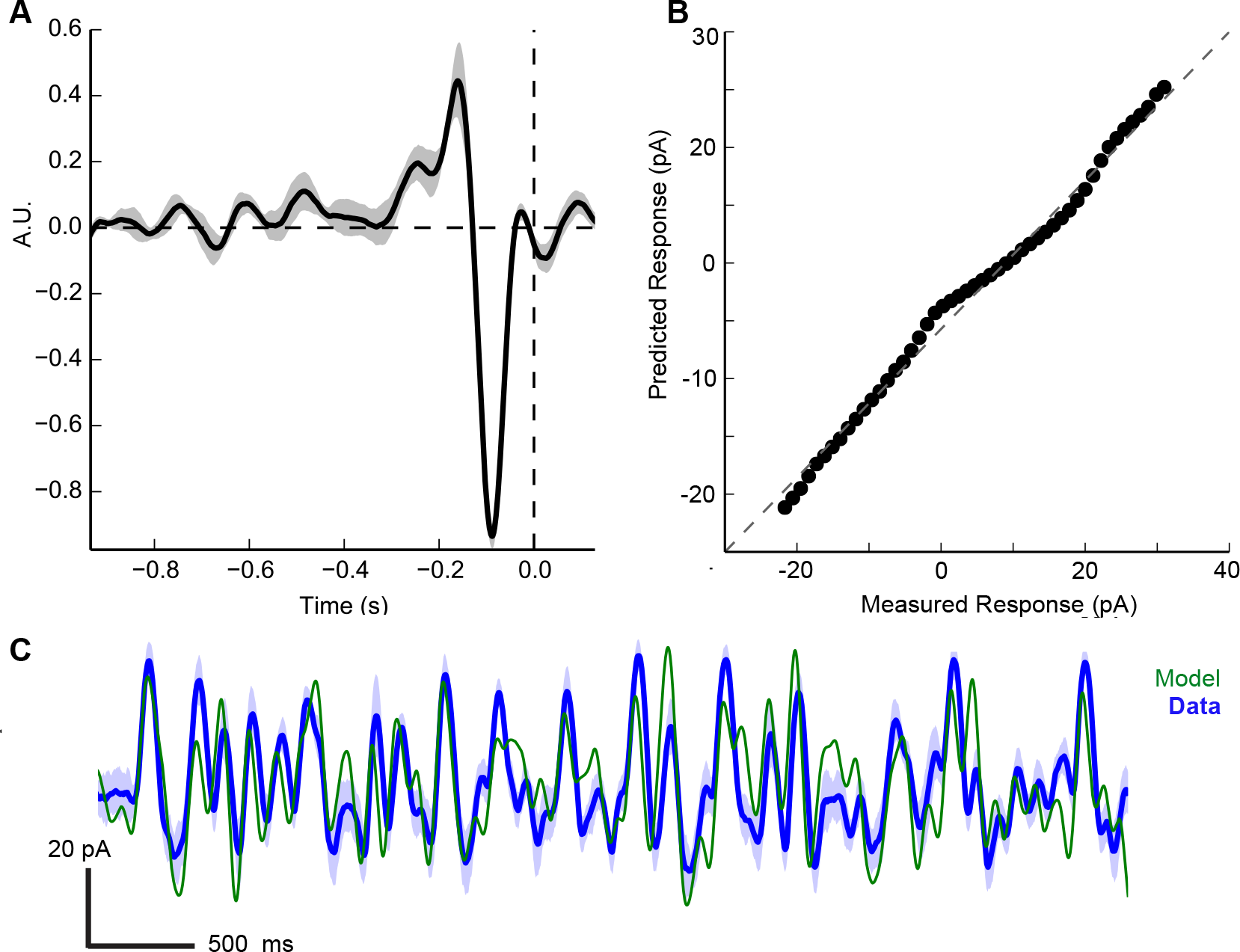
LN model of ON bipolar cell current. **A.** Linear filter of a voltage-clamped ON bipolar cell stimulated with 33% contrast binary white noise. **B.** Static nonlinearity calculated by comparing the measured response with the convolution of the stimulus and linear filter. **C.** LN model prediction (green trace) vs. average measured current (blue trace). Light blue trace represents standard deviation (5 trials).

The LN model assumes that both the linear filter and nonlinearity are time-invariant. To test this assumption, we presented paired 50 ms flashes at 100% contrast while varying the inter-flash interval (IFI) from 0 to 400 ms (Figure 5A). The principle of linear superposition predicts that the measured response should equal the sum of the response to two individual flashes with the proper time delay. A cell that exhibits history dependence, on the other hand, would fail this test for linear superposition because the first flash would change the response sensitivity of the cell for the second flash.

**Figure 5.**
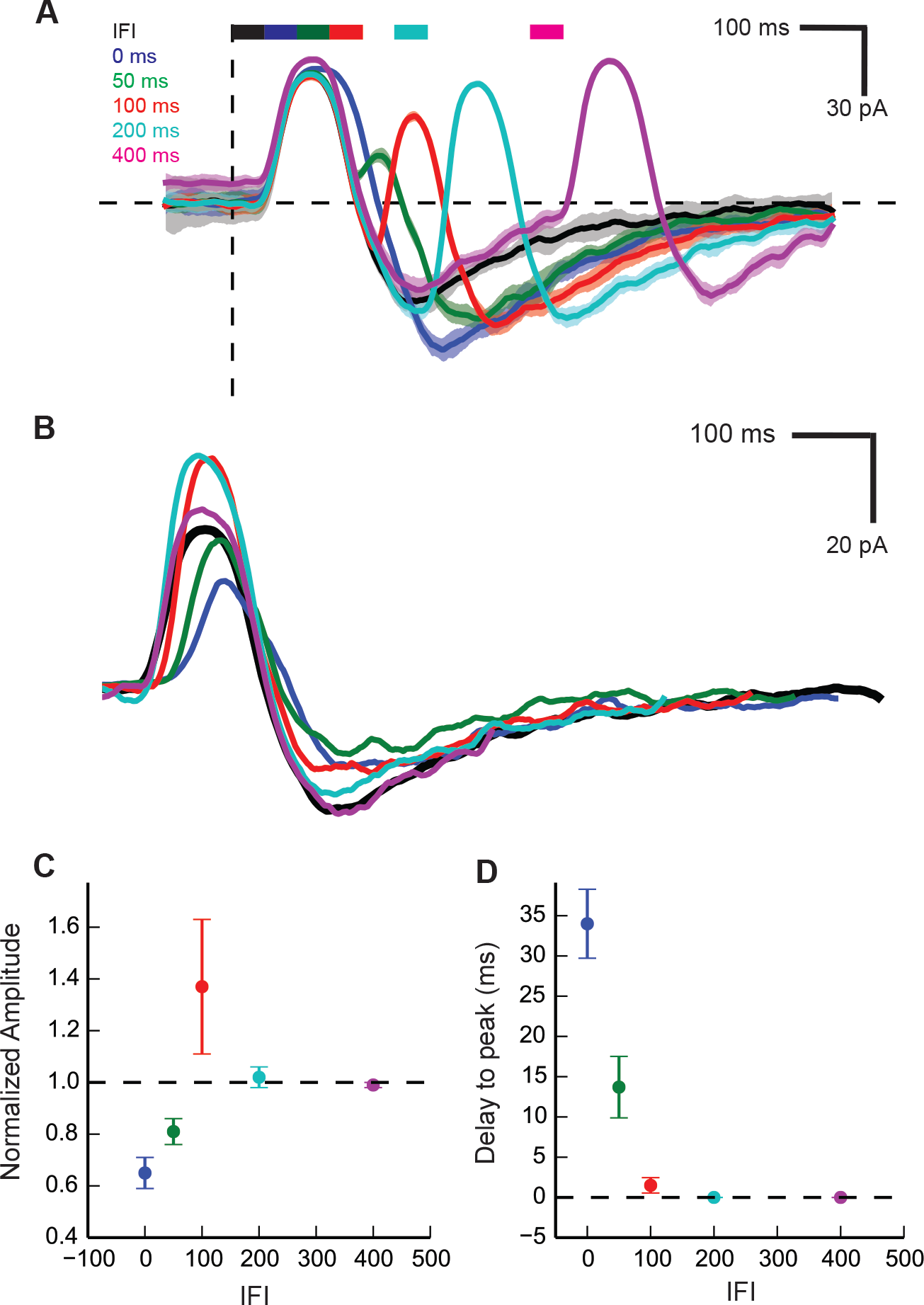
Paired high contrast flashes reveal gain control in ON bipolar cells. **A.** Responses to paired flashes with varying inter flash intervals (IFI) plotted against a single flash (black) for reference. Shaded regions around each trace represent standard deviation. Each flash was 50 ms in duration and 100% contrast. Vertical and horizontal dashed lines represent time zero and zero current, respectively. **B.** Single flash response (black) compared with the residuals after subtracting the single flash from paired flashes for all IFIs. **C.** Normalized amplitude of peak outward current for all IFIs. Error bars indicate SEM (n=9). **D.** Time delay to peak outward current for all IFIs. Error bars indicate SEM (n=9).

To test this prediction, we examined the residual after subtracting the single flash from the response to the paired flash, as shown in Figure 5B. The outward current exhibited a delay to peak for the 0 ms and 50 ms IFI conditions (the former case is identical to a single 100 ms duration flash) as well as a modulation of the peak amplitude that varied as a function of IFI. Additionally, the amplitude of the outward current never exceeded the amplitude of the single flash, which is consistent with the saturation effect described in Figure 3. As shown in Figure 5C, the reduction in peak amplitude was significant for 0 ms and 50 ms IFI (p=0.001 and p=0.014, respectively, n=9). The delay to peak, quantified for the population of cells in Figure 5D, was significant for the 0 ms and 50 ms IFI (p=0.0005 and p=0.019, respectively). By 400 ms, both the time course and amplitude of the second flash response matched the single flash response, indicating recovery to the baseline state before stimulation. This paired flash depression and subsequent recovery indicate the presence of a rapid, transient gain control at the input to ON bipolar cells.

In order to quantitatively describe the transient input nonlinearities present in ON bipolar input current, we constructed a phenomenological model of the light to ON bipolar current system. A block diagram of this model is shown in Figure 6A. The transformation from light to photoreceptor voltage can be modeled as a linear system, even at high contrasts [38]. To calculate the voltage response at the photoreceptor, *L(t),* we convolved the light stimulus (in units of Weber contrast) with an alpha function that approximated the impulse response of the photoreceptor membrane. The amount of glutamate released at the cone pedicle as a function of membrane voltage goes to zero in response to very strong bright flashes. We approximate this asymmetric synaptic activation by passing the voltage response *L(t)* through a static saturating nonlinearity *N*_*1*_*(t)* that amplifies the response to dark flashes and suppresses the response to dark flashes to yield the glutamate release, *u(t)*. This nonlinearity is similar to the synaptic activation described in [39], in which the resting voltage of the photoreceptor sits at the foot of the sigmoidal synaptic activation curve.

In the next stage of the model, we sought to capture the dynamics of the mGluR6 receptor to TRP channel system. The G-protein coupled signaling cascade in ON bipolar cells has not been fully characterized, ruling out the possibility of producing a biophysically realistic model. The LNK model published by [37] can accurately recapitulate the rapid transient voltage response of ganglion cells at transitions from low to high contrast by incorporating first-order rate dynamics. By adding a similar kinetics block in our model, we can simplify the signaling elements of the mGluR6 second messenger cascade into three pools of components: a ready pool *R* that determines the instantaneous gain of the system, an activated pool *A* that gates the TRP channel, and an inactivated pool *X* that must be converted back to the ready state to participate in signaling. This form of model produces gain control as well as an overshoot at stimulus offset, and does not depend on a particular biophysical mechanism.

**Figure 6.**
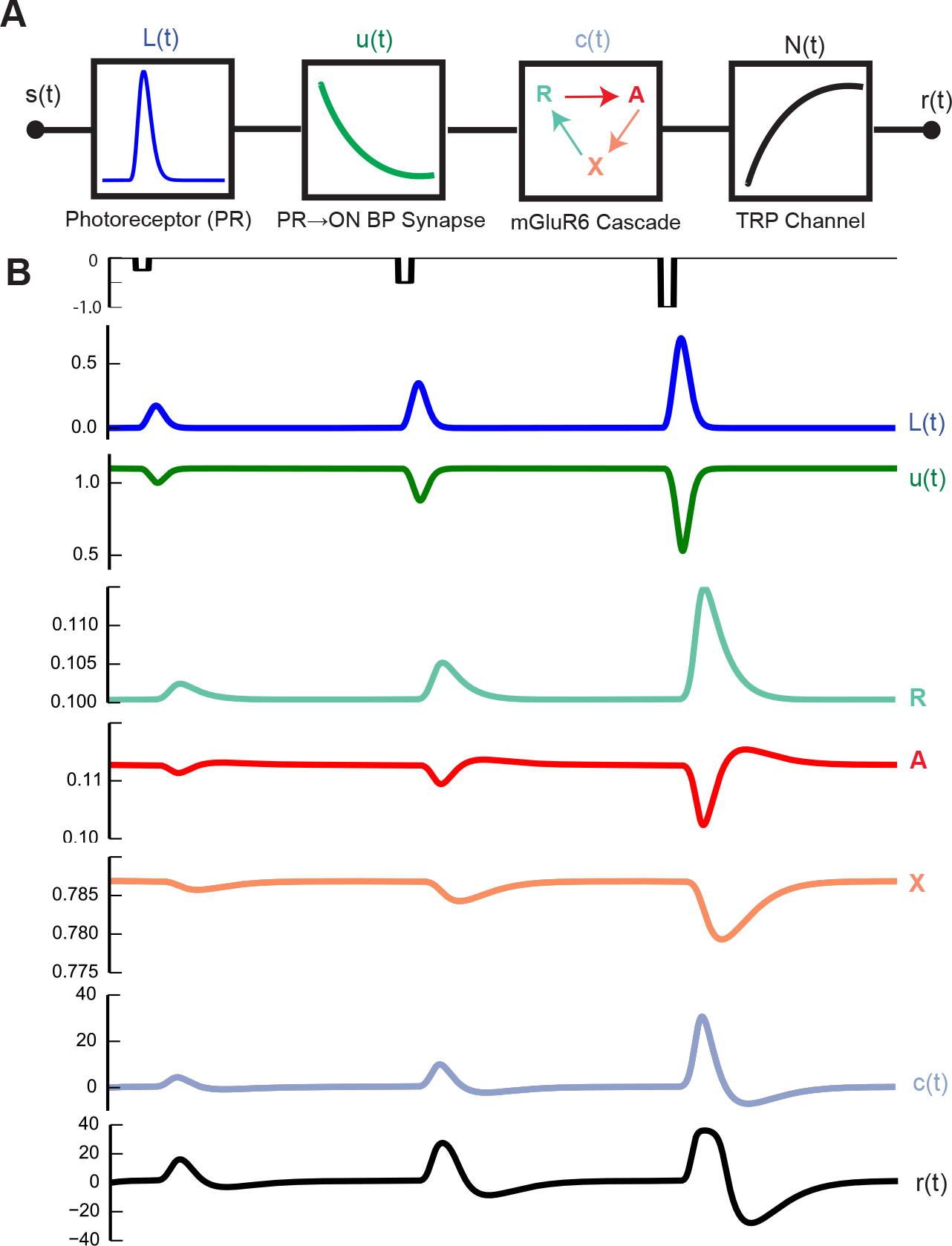
LNKN model of light to ON bipolar cell current. **A.** Block diagram of the model. **B.** Dynamics of each internal state variable of the LNKN model during responses to dark flashes of three different contrasts.

The maximum outward current of the system saturates at high contrast and long duration (see Figure 3). To model this second rectification, the current response *I(t)* is calculated by passing the output of the kinetic block *A(t)* through a saturating nonlinearity *N*_*2*_(*t*) that limits the maximum amplitude and scales the response to units of pA. The internal variables of this model are shown for dark flashes of increasing contrast in Figure 6B.

This linear-nonlinear-kinetic-nonlinear (LNKN) model successfully captures many of the key features of the ON bipolar response. In response to 33% white noise, the normalized root mean squared error between model and data is 13.4%, compared to 18% for the LN model (Figure 7A vs. Figure 4). The response to a single flash is biphasic, with a significant overshoot at higher contrasts. The maximal inward current and outward current both saturate at higher contrast, and the amplitude of the response to dark flashes is larger than the response to a bright flash of equal contrast (Figure 7B and 7C), recapitulating the asymmetry shown in Figure 1 and the rectification described in Figures 2 and 3.

**Figure 7.**
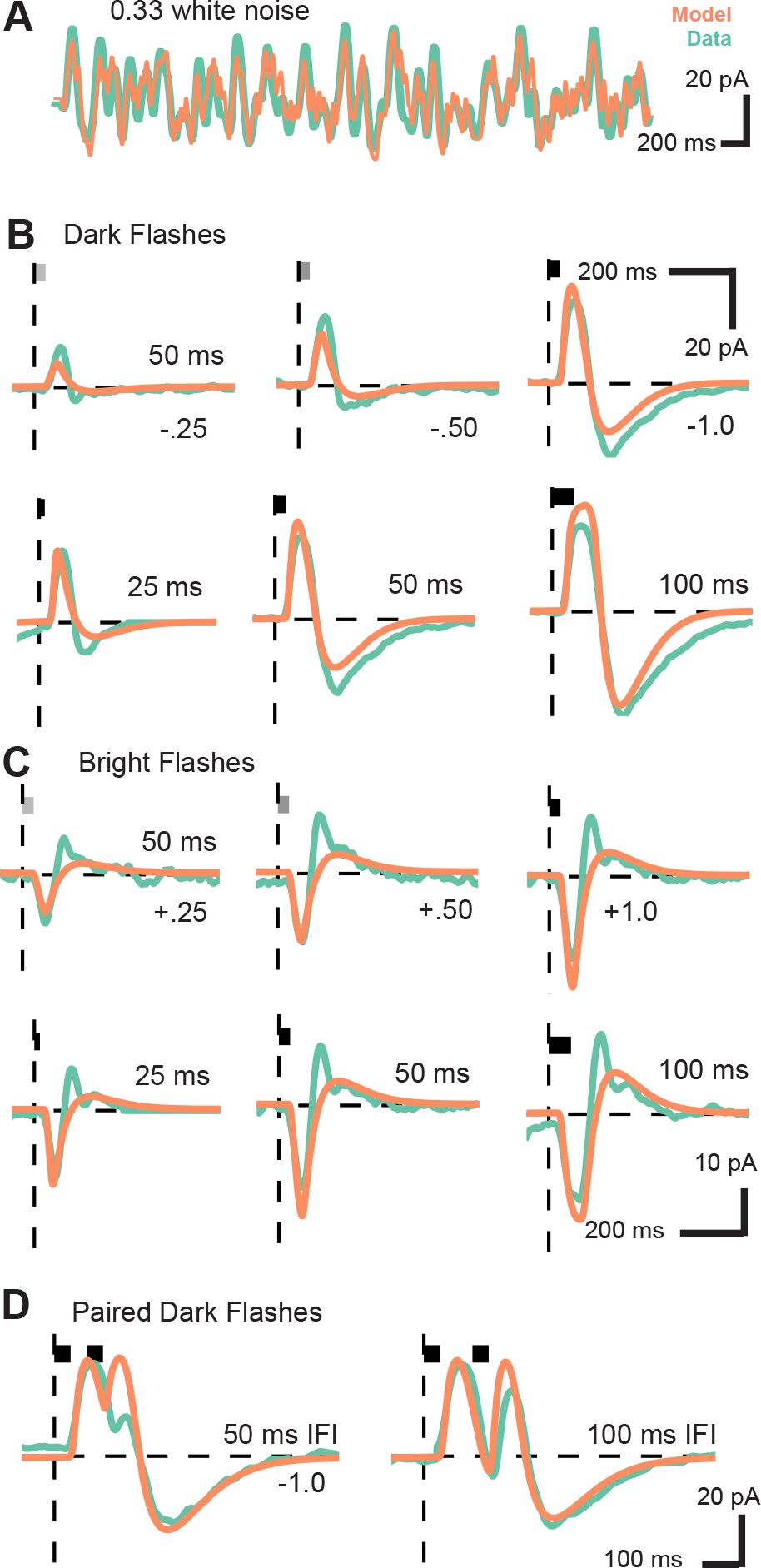
The LNKN model captures the key features of ON bipolar cell responses. **A.** Model (orange) and data (green) in response to 33% white noise. **B.** Model (orange) and data (green) in response to dark flashes of 25, 50 and 100 ms duration and dark flashes of 25%, 50% and 100% contrast. **C.** Model (orange) and data (green) in response to bright flashes of 25, 50 and 100 ms duration and bright flashes of 25%, 50% and 100% contrast. **D.** Model (orange) and data (green) in response to paired flashes with 50 ms and 100 ms inter-flash interval.

A key feature of the model is the emergence of the large overshoot at −100% contrast. The rate of *X*→*R* is ~10x slower than the rate of *R*→*A* and *A*→*X*. At stimulus offset, the inactive pool *X* remains higher than baseline, reducing the ready pool *R* such that the activation *A* is reduced below the baseline level. This produces the overshoot at the offset. Before the final rectification stage *N*_*2*_*(t),* the amplitude of the peak model response to a 100% flash is approximately twice the response to a 50% contrast flash. The rectification suppresses the amplitude of the outward component while preserving the large inward overshoot generated by the kinetic block.

## Discussion

Our study describes three distinct types of nonlinearity in ON bipolar cells. The first is the asymmetry between the response to bright and dark flashes at high contrast due to the suppression of the bright flash response (Figure 1). The second is the saturation of the OFF response revealed by the failure of the linear superposition model (Figure 3). The third is the rapid, transient paired flash depression (Figure 5).

### Mechanisms of rectification

At low contrast, ON bipolar cells operate in the linear regime, with symmetric responses to increments and decrements. However, at high contrast, both bright and dark flashes appear to exceed the dynamic range of the mGluR6 pathway (Figure 2). Fahey and Burkhardt reported that in ON bipolar cells, the maximum amplitude of the voltage response to negative contrast steps grew as a function of background intensity, while the response to positive steps remained constant [40]. The asymmetry observed in Figure 1, in which the OFF response of ON bipolar cells is significantly larger than the ON response, was measured with a background intensity well into the photopic range, and is thus consistent with these findings. Furthermore, our measurements were performed in voltage-clamp, which eliminates the transformation from synaptic current to membrane voltage as a source of this asymmetry. One plausible mechanism is that a sufficiently intense bright flash hyperpolarizes cone photoreceptors to the point where they no longer release glutamate, limiting the amplitude of the inward current produced. In our model, this rectification is modeled by the synaptic activation function *u(t)*. Alternatively, a strong bright flash could sufficiently deactivate the mGluR6 pathway to open all of the available TRPM channels, limiting the maximal conductance.

The saturation of the OFF response to longer duration flashes (Figure 3), on the other hand, could result from a number of possible mechanisms. At high contrast, all of the readily releasable pool at the cone ribbon synapse could be released at once. Alternatively, the flooding of the synaptic cleft with glutamate could saturate all available mGluR6 receptors. A third possibility is that sufficiently strong glutamate input could drive the mGluR6 cascade to close all of the available TRPM channels. All three mechanisms ultimately limit the maximum amplitude of the response and are captured by the *N*_*1*_*(t)* rectification in the model.

### Mechanisms of rapid gain control

The LN models calculated to study contrast adaptation in ON bipolar cells [28] and the transfer functions calculated with sinusoidal stimulation [41] represent the steady-state response properties of the cell. However, these models were constructed by explicitly excluding transient responses at the start of the sequence, limiting their utility in predicting the response to transient stimuli. The ON bipolar cells recorded in this study were approximately linear at steady-state and low contrast stimulation (Figure 4), but exhibited a time-dependent modulation of gain in response to 100% contrast dark flash stimuli (Figure 5A). Sharp gain reduction occurred within 50 ms of the first flash, ruling out slower gain control mechanisms such as the adaptation to mean luminance, which has a timescale of seconds, and calcium-mediated desensitization of the mGluR6 cascade [25], which has a timescale of hundreds of milliseconds. Furthermore, the reduction in gain occurred under voltage-clamp, ruling out the possibility of negative feedback via voltage-gated ion channels.

One possible mechanism is feedback from amacrine cells onto the bipolar cell terminals. Bipolar cells are electrotonically compact, so inhibitory input at the axon terminal forms a component of the measured current. Another possibility is that the gain reduction is mediated by negative feedback within the mGluR6 transduction cascade. In addition to intracellular calcium, there are many elements of the second messenger cascade that could contribute to the temporal response properties observed in the input current. For example, in rod bipolar cells, glutamate binds to mGluR6, activating the alpha subunit of the G-protein, which in turn triggers DAG→PLC→PKCα. The activation of PKCα has been shown to potentiate the current through the TRPM1 channel [42]. While this exact pathway does not exist in ON cone bipolar cells, perhaps there is a modulatory signaling element within the mGluR6 cascade that can modulate the gain further than the 20% gain reduction explained by the 3-state kinetic block.

### Limitations of the model

While the model successfully captures many of the salient response features of the measured input current, it does not capture the rapid and transient outward current following bright flashes (Figure 7C). One possibility is that this outward current is caused by inhibitory feedback from an amacrine cell. Such a process would be completely outside of the scope of the model we have formulated. In addition, the 3-state kinetics block can only produce a ~20% reduction in gain. This gain reduction is not sufficient to match the >20% gain reduction of a paired second flash that occurs 50 ms after the first flash. The model accurately predicts the response for the 100 ms inter-flash interval (Figure 7D). In general, our model aims to achieve a balance between accuracy and simplicity. Simplicity is valuable because the resulting model has relatively few parameters that can be constrained by existing data. As further information becomes available about the biochemical cascade initiated by the activation of the mGluR6 receptor as well as about the types of amacrine cells that provide input to ON bipolar cells, further refinements of this form of model will likely become possible.

### Implications for visual coding

In general, rectification and gain control mechanisms serve to compress the signals to maximize efficient use of the dynamic range of neuronal output. While the distribution of contrasts in natural scenes is heavily weighted towards small contrasts and low intensities, the distribution has a long tail of high contrast stimuli [43–46]. Thus, an important operation for the retina to carry out is to compress this high dynamic range of light intensities into the limited dynamic range of neural signaling. We observed significant nonlinearities in the input current to ON bipolar cells starting at Weber contrasts of ±50%. Given the wide range of light intensities found in natural scenes, we expect that this nonlinear signaling regime will be achieved often. We also observed that nonlinear signaling in ON bipolars was enhanced by changes in light intensity that persisted for 50 ms or longer. Naturalistic visual stimuli also exhibit a wide range of durations over which light intensity fluctuates [43], which again makes it likely that nonlinear signaling in ON bipolars will be engaged.

Another important feature of natural visual scenes is the overrepresentation of dark contrasts [47]. This property results from the skew of the distribution of light intensities and has been related to the fact that retinas of many species have a higher density of OFF-type retinal ganglion cells than ON-type [48–51]. In this vein, we observed a much higher gain for OFF-responses than for ON – both in the peak outward current and in the overshoot inward current. This higher gain is matched to the greater prevalence of dark contrasts and may help to better encode the spatial information contained in dark contrasts.

